# NeuroFreq: a MATLAB Toolbox for Time-Frequency Analysis of M/EEG Data

**DOI:** 10.1101/2023.11.01.565154

**Authors:** Eric Rawls

## Abstract

Time-frequency (TF) analysis of M/EEG data enables rich understanding of cortical dynamics underlying cognition, health, and disease. There are many algorithms for time-frequency decomposition of M/EEG neural data, but they are implemented in an inconsistent manner and most existing toolboxes either 1) contain only one or a few transforms, or 2) are not adapted to analyze multichannel, multitrial M/EEG data. This makes entry into time-frequency daunting for new practitioners and limits the ability of the community to flexibly compare the performance of multiple TF methods on M/EEG data. This paper introduces the **NeuroFreq** toolbox for MATLAB, which includes multiple TF transformation algorithms that are implemented in a consistent fashion and produce consistent output. The toolbox includes TF decomposition algorithms of both linear and quadratic classes, utilities for resampling, averaging, and baseline correction of TF representations, and tools for visualizing and interacting with single-trial or averaged TF representations over multiple channels. This paper introduces these utilities, and applies them to synthetic and EEG data to demonstrate the **NeuroFreq** toolbox.

## 1. Time-Frequency Analysis of M/EEG Data

Time-frequency (TF) analysis is a vital tool for interpreting brain electrophysiological data including electroencephalography (EEG) and magnetoencephalography (MEG). Neural oscillations, rhythmic fluctuations in electrical potentials, facilitate information transfer across distributed networks in the brain (Buzsáki and Draguhn 2004; Buzsaki 2006) and play a critical role in cognition (Ward 2003). The insights offered by TF analysis are tempered by practical challenges and inconsistencies in the application of these methods. There are many algorithms for TF decomposition with differing benefits and drawbacks. These benefits or drawbacks can be difficult to assess or quantify, since different TF algorithms are implemented inconsistently across multiple software packages. The following briefly describes the major classes of TF analysis algorithms, followed by a discussion of the limitations of existing software tools for TF analysis of M/EEG data structures. These limitations justify the development of the **NeuroFreq** Toolbox for time-frequency analysis of M/EEG data.

### 1.1. Classes of TF Decomposition Algorithm

TF algorithms can be broadly categorized into linear and quadratic approaches. Linear TF methods are typified by the short-time Fourier transform (Gabor 1946), which is characterized by an inflexible time and frequency resolution tradeoff resulting from using fixed time windows, and thus can provide high time or frequency resolution but usually not both simultaneously. This problem is addressed by methods like wavelet convolution (Grossmann and Morlet 1984) that can provide enhanced frequency resolution at the cost of decreased temporal resolution for low frequencies, and vice-versa for high frequencies. In contrast, quadratic TF techniques, including particularly the “Reduced Interference Distributions” (RID) (Jeong and Williams 1992) which are in Cohen’s class of TF distribution (Cohen 1995), are sometimes sought in M/EEG analysis for their uniform high resolution in both time and frequency. The application of RID, particularly as employed in the ERP-TF-PCA method (Bernat, Williams, and Gehring 2005), represents an example of this high-fidelity approach to TF analysis. The different TF algorithms are described in Section 3. See Figure 1 for an illustration of the time-frequency tradeoff inherent in different TF algorithms. Interestingly, a direct comparison between the results of linear and quadratic TF transforms is often difficult because most available toolboxes only provide access to a limited number of TF algorithms. These available toolboxes, and the algorithms they provide access to, are briefly summarized in the following section.

**Figure 1.**
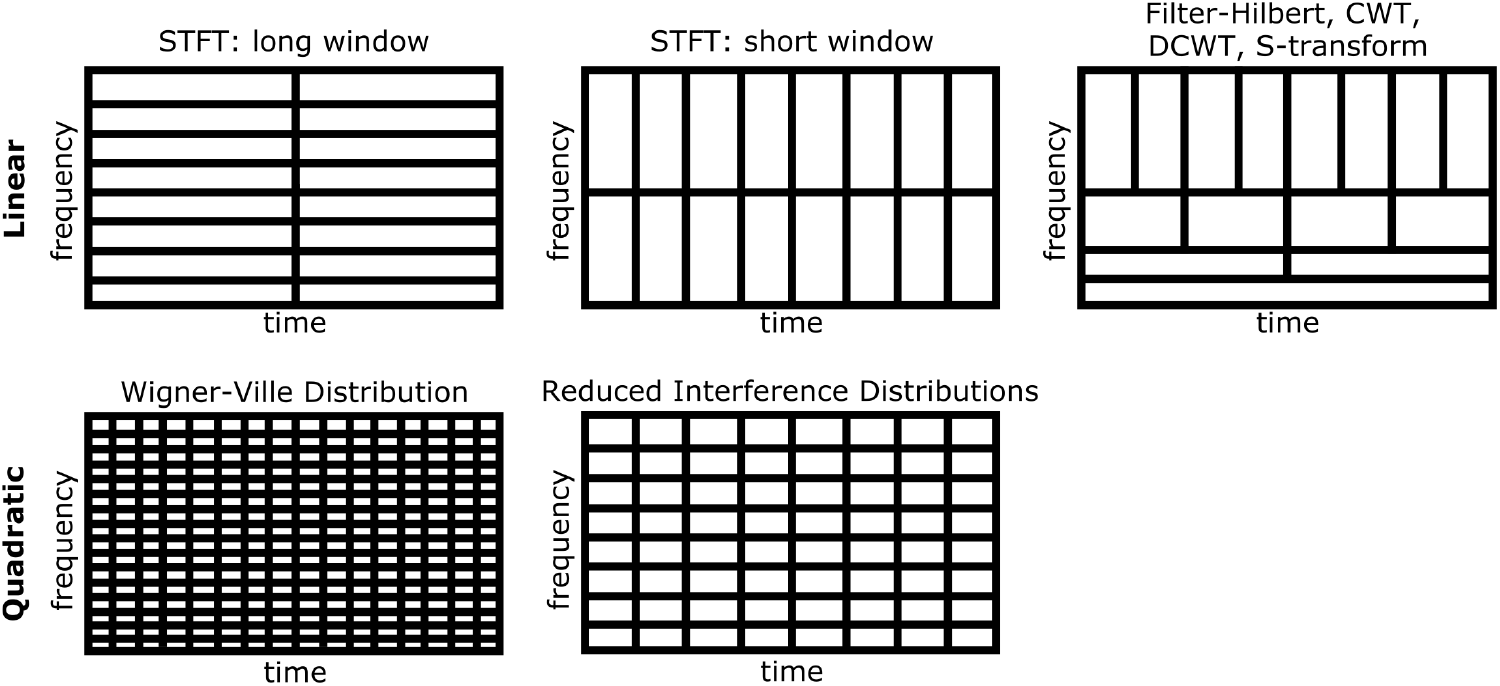
Time-frequency resolution tradeoff inherent in different TF algorithms. STFT uses fixed time windows leading to an inflexible TF tradeoff. CWT, DCWT, and S-transform have increased temporal resolution at higher frequencies due to compression of a Gaussian function; the same occurs for filter-Hilbert as high-frequency filters have decreasing numbers of points. The Wigner-Ville distribution has excellent joint time-frequency resolution, but suffers from cross-terms that render it unusable for most practical applications. The Reduced Interference Distributions solve this problem by applying kernel functions to the Wigner-Ville distribution, significantly masking cross-terms at the expense of slightly lower TF resolution.

### 1.2. Previous MATLAB Toolboxes for M/EEG TF Analysis

TF analysis of M/EEG data has seen advancements through various computational toolboxes but limitations persist across available tools. **EEGLAB** (Delorme and Makeig 2004) offers a common platform for TF analysis but largely confines its methods to wavelet transform and STFT. **FieldTrip** (Oostenveld, Fries, Maris, and Schoffelen 2010) emphasizes wavelet and STFT methods. **BrainStorm** (Tadel, Baillet, Mosher, Pantazis, and Leahy 2011) includes a Hilbert transform-based method in addition to the STFT and the Morlet wavelet transform. **ERPWaveLab** (Mørup, Hansen, and Arnfred 2007) also provides only STFT and wavelet-based TF decomposition. The **Psychophysiology Toolbox** (PTB; http://www.ccnlab.umd.edu/Psychophysiology_Toolbox/) is an exception by including quadratic RID methods, but it also provides a restrictive set of methods by including only continuous wavelet transform and RIDs. The shared limitation across these toolboxes lies in their confinement to one or a few TF algorithms, hindering comprehensive comparisons and the ability to select the most suitable method for a given research question. A practical consequence of this limitation is that the results of different time-frequency algorithms are in different formats, making direct comparison among methods difficult. These limitations necessitate the development of a new tool for TF analysis for M/EEG data: the **NeuroFreq** Toolbox for MATLAB.

**Table 1:** Comparison of existing software for time-frequency analysis of EEG/MEG data. (✓) denotes that the software (row) implements the specific method (column).

|  | STFT | Filter-Hilbert | Demodulation | CWT | Morlet Wavelet | S-Transform | Binomial RID | Born-Jordan RID | RID-Rihaczek |
| --- | --- | --- | --- | --- | --- | --- | --- | --- | --- |
| EEGLAB | ✓ |  |  |  | ✓ |  |  |  |  |
| FieldTrip | ✓ |  |  |  | ✓ |  |  |  |  |
| BrainStorm | ✓ | ✓ |  |  | ✓ |  |  |  |  |
| ERPWaveLab | ✓ |  |  |  | ✓ |  |  |  |  |
| PTB |  |  |  | ✓ |  |  | ✓ | ✓ | ✓ |
| BESA |  |  | ✓ |  |  |  |  |  |  |
| NeuroFreq | ✓ | ✓ | ✓ | ✓ | ✓ | ✓ | ✓ | ✓ | ✓ |

### 1.3. Brief Outline of the Paper

The remainder of the paper is structured as follows. Section 2 describes the contents of the **NeuroFreq** Toolbox. First the graphic interface and the high-level scripting utilities, including functions for data preparation and TF transformation, are introduced. Also described are optional post-processing utilities for TF-transformed signals, and utilities for interacting with and visualizing TF-transformed data. Section 3 introduces the included TF transformation algorithms, including their mathematical basis and the functions used to apply them in **NeuroFreq**. Section 4 demonstrates the capabilities of the **NeuroFreq** toolbox by applying the included TF transforms to data. TF algorithms are applied to simulated data and to sample EEG data from a visual experiment. Section 5 concludes the paper.

## 2. NeuroFreq Toolbox Contents

### 2.1. Overview

**NeuroFreq** is a MATLAB toolbox for flexible time-frequency analysis of M/EEG data. **NeuroFreq** includes a greater variety of time-frequency transformation algorithms than existing toolboxes, and is flexible with the capability to run both linear and quadratic time-frequency transformations. All algorithms accept as input either **EEGLAB**-formatted (.set) datasets or data of channels X times X trials, where both channel and trial dimensions are optional. All functions are suitable for real or complex input. All output are formatted the same, enabling easy direct comparisons between different algorithms. **NeuroFreq** is not meant to replace existing packages for preprocessing M/EEG data or for artifact removal, and users should preprocess their data using an existing software environment prior to import into **NeuroFreq** for analysis. **NeuroFreq** functions are compatible with EEGLAB .set files, and users can directly import EEGLAB files into **NeuroFreq** using the GUI or apply TF transforms directly to EEGLAB-formatted datasets.

### 2.2. Data Preparation

The **NeuroFreq** Toolbox includes a utility to prepare already preprocessed artifact-free time-domain signals for TF transformation. This function does not clean or preprocess M/EEG data. Instead, the function 1) mean centers each sensor/epoch of data, 2) removes quadratic trends, 3) applies a cosine-square taper to the beginning and final 5% of datapoints, and 4) makes real signals analytic using the Hilbert transform. These operations are done for each channel and trial of data. This preparation reduces spectral misspecification due to edge effects and quadratic trends. The function is called as follows:

~~~
>> EEG = nf_prepdata(EEG, center, detrend, taper, analytic);
~~~

Where EEG can be either 1) an **EEGLAB**-formatted (.set) data structure or 2) a data matrix of dimensions channels X time points X trials. The arguments center, detrend, taper, and analytic should all be 0 or 1, indicating whether to do the specified operation. The prepared data is returned in the .data field of the **EEGLAB** .set if an **EEGLAB** .set is the input, and the prepared data is returned in a matrix if a matrix is the input.

### 2.3. TF Decomposition

The primary functionality of the **NeuroFreq** toolbox is to perform efficient, high-quality TF decompositions with a wide variety of algorithms. Following optional data preparation, M/EEG time series data can be transformed to a TF representation using any of the included algorithms, including several implementations of linear and quadratic decomposition methods as specified below. One main feature of the **NeuroFreq** Toolbox is the inclusion of a high-level function for TF computation, which provides access to every TF algorithm using a unified syntax and a single function with the option to specify the method of choice. This function accepts keyword-argument pairs that differ depending on the algorithm selected, and uses sensible defaults in the case of unspecified keyword-argument pairs. The high-level function facilitates adoption of the individual TF decomposition methods by accepting as input an EEGLAB-formatted (.set) data structure, which is among the most common software packages for preprocessing M/EEG data. The high-level function is called as follows:

~~~
>> TF = nf_tftransform(EEG, ‘method ‘, ‘methodarg ‘, ‘key1 ‘, ‘arg1 ‘, …);
~~~

Where 1) EEG is an **EEGLAB**-formatted (.set) data structure, and 2) ‘methodarg ‘ can be any of the transforms detailed in Section 3 of this paper. Additional keyword-argument pairs differ by algorithm and are detailed extensively in the documentation of `nf_tftransform.m`. Section 3 introduces the theoretical background for each algorithm included in **NeuroFreq**.

### 2.4. Post-Processing of TF Transformed Data

While TF computation is the primary goal of **NeuroFreq**, the toolbox also includes a small number of utilities aimed at post-processing TF data for various experimental purposes. These utilities include 1) a function for trial-level averaging and baseline correction of TF data, 2) a function for downsampling TF transformed data in time and frequency, and 3) a function for aggregating multiple TF-transformed datasets into a multi-subject dataset.

The averaging function `nf_avebase.m` averages together single trials of TF data prior to applying a baseline correction using either decibel, percent change, z-score, or no correction. For TF decompositions that return phase estimates, this function averages phase over trials to estimate inter-trial phase coherence (ITPC). This function is called as:

~~~
>> TF = nf_avebase(TF, ‘blmethod ‘, bltimes, trlvec);
~~~

Where ‘blmethod ‘ can be ‘dB ‘, ‘percent ‘, ‘zscore ‘, or ‘none ‘ and bltimes includes the time samples to be included in the baseline. trlvec is a vector of the same length as the number of trials in the data, containing unique identifiers referring to trials that should be averaged together. If ‘blmethod ‘ is left out then the function defaults to ‘percent ‘ correction, if bltimes is left out the function defaults to all times before 0 (event onset), and if ‘trlvec ‘ is left out the function defaults to averaging all trials together.

The resampling function `nf_resample.m` can resample TF representations to new time or frequency vectors using linear interpolation. This function is useful for TF methods that produce a redundant frequency sampling, such as the S-transform or CWT, and can also be used to reduce the number of time or frequency points in a representation in order to reduce the number of statistical tests carried out at a later step. This function is called as:

>> TF = nf_resample(TF, tVec, fVec);

Where tVec contains the new times to resample to and fVec contains the new frequencies to resample to. Either tVec or fVec can be left blank, in which case only one dimension of the TF representation is resampled.

The aggregation function `nf_aggregate.m` opens a menu for selecting multiple subject-level TF datasets. If the datasets are averaged and contain consistent data sizes, they will be conacatenated into a multi-subject dataset suitable for inspecting the results of entire studies at once. This function is called as:

~~~
>> TF = nf_aggregate();
~~~

No arguments are required since the function opens a menu for selecting datasets for aggregation.

### 2.5. Visualization and Inspection

Following TF decomposition, it can be helpful to visualize the TF matrices as surfaces or scalp topographic plots. For single-trial data this can help visually identify remaining artifacts in the data that should be deleted, and for all TF transformations this can help clarify scalp distributions and temporal characteristics of power. A simple plot of TF power, averaged over all trials and channels, can be shown with the function

~~~
>> TF = nf_tfplot(TF);
~~~

And an interactive visualization of TF structures, including topographic plots and surface plots (at specified sensors) can be generated with the function

~~~
>> TF = nf_viewerapp(TF);
~~~

The viewer app allows the option of scrolling through single-trials for unaveraged data, and scrolling through time and frequency points for averaged or single-trial data. The topographic plot will automatically update when the trial, time point, or frequency point are changed, and the surface plot will automatically change when the scalp electrode is changed. This function uses code included in Mike X. Cohen’s excellent open-source tool `tfviewerx.m` (Cohen 2014), but includes extra utilities for visualizing and scrolling single-trial data.

## 3. Time-Frequency Algorithms

### 3.1. Notation for Time-Frequency Decompositions

Throughout the following, let *t* represent time and *x*(*t*) represent the time-domain signal, *f* represent frequency, *w*(*t*) represent a window function, *τ* represent a time shift or lag, ^*\**^ represent the complex conjugate, and *ϕ*(*t, f*) denote the time-frequency representation of *x*(*t*). *σ* additionally represents a frequency shift, alternately called doppler, which is only used in the quadratic transforms. All integrals are taken from negative to positive infinity.

### 3.2. Linear Time-Frequency Decomposition Methods

Linear time-frequency methods offer a direct and efficient approach to time-frequency analysis. They provide a clear and computationally efficient representation of the signal’s time-frequency content but often exhibit a fixed or rigid trade-off between time and frequency resolution. These methods often provide high frequency resolution at the expense of temporal resolution, or vice versa. Thus they are best suited for situations where such a trade-off is acceptable.

#### Short-time Fourier Transform

The short-time Fourier Transform (STFT) is the most established method for time-frequency analysis (Gabor 1946). The STFT cuts data into windows of a specified length and with a specified percentage overlap. The data segments are then windowed and a discrete Fourier transform is computed on each data segment. This estimates the Fourier spectrum for the center point of each data segment. The STFT is mathematically defined as

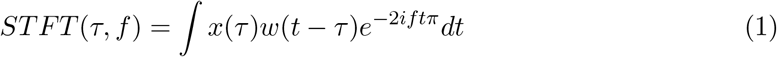

indicating that the time domain signal *x*(*t*) is first multiplied by a window function *w* centered at time *t*, followed by computing the discrete Fourier transform. Customarily *t* = *τ*.

The benefits of the STFT are: 1) it produces a high signal-to-noise ratio, which makes the STFT robust to noise; 2) it is efficient to compute; 3) almost every programming language has built-in support for calculating STFT. The primary disadvantage of the STFT is that it uses fixed window lengths (Cohen 1995). The limitation of constant time-frequency resolution can pose challenges when trying to investigate phenomena occurring in different frequencies in M/EEG data, where it can be important to have high frequency resolution at lower frequencies and high temporal resolution at higher frequencies (Cohen 2014). The STFT’s constant window length assumes stationarity within the window, which may not be a valid assumption for neurophysiological data (Kaplan, Fingelkurts, Fingelkurts, Borisov, and Darkhovsky 2005). This can result in distortion or leakage effects, reducing the accuracy of time-frequency representations. The STFT is calculated using

~~~
>> TF = nf_stft(data, Fs, window, overlap, fRes, plt);
~~~

Where data is a 1/2/3D tensor of dimensions channels X time X trials, Fs is the sampling rate of the data in Hz, window is the length of the window in seconds, overlap is the percentage of overlap, fRes is the requested frequency resolution, and plt is 0 or 1 indicating whether or not the user would like a summary plot to be produced following transformation.

#### Filter-Hilbert

The filter-Hilbert method combines bandpass filtering with Hilbert transformation. The filter-Hilbert method doesn ‘t impose a predefined kernel shape, allowing for a custom design of bandpass filters based on the specific needs of the analysis. The filter-Hilbert operation be described by:

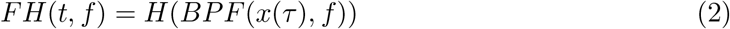

Where *H* represents the Hilbert transform and *BPF* represents a bandpass filter.

While the filter-Hilbert method can offer excellent frequency specificity depending on filter design, the method also has some potential drawbacks. The need to select a filter order and bandwidth introduces additional complexity to the analysis. The filter order determines the sharpness of the transition between the passband and the stopband of the filter, with higher orders resulting in steeper roll-off but potentially causing more distortion due to the longer filter impulse response (Cohen 2014; Luck 2014). Similarly, the bandwidth selection can impact the resolution in time and frequency.

Our implementation of the filter-Hilbert method filters M/EEG signals into specified frequency bands using Butterworth filters (Butterworth 1930). Users may select the order of the filter and the bandwidth of the filter. Butterworth filters are known for their frequency response with no ripple, making them a popular choice in many signal processing applications. The filter-Hilbert transform is calculated using

~~~
>> TF = nf_filterhilbert(data, Fs, freqs, fBandWidth, order, plt);
~~~

Where data is a 1/2/3D tensor of dimensions channels X time X trials, Fs is the sampling rate of the data in Hz, freqs is a vector of center frequencies for filtering, fBandWidth is the frequency bandwidth of the filters, order is the order of the filters, and plt is 0 or 1 indicating whether or not the user would like a summary plot to be produced following transformation. fBandWidth can be specified as a single element (e.g. 1) in which case all filters will use the same width, or as a two-element vector (e.g. [1 6]) in which case the bandwidth of filters will be 1 Hz at the lowest frequency and will linearly increase to 6 Hz at the highest frequency.

#### Complex Demodulation

Complex Demodulation calculates time-frequency representations by multiplying the real-valued signal by a complex sine wave of a specific frequency, which demodulates the power at that frequency so that it is concentrated at low frequencies. Then, the signal is lowpass filtered in order to recover power and phase at the frequency of the sine wave. While this method originated in radio signal processing and is uncommonly applied to M/EEG recordings, complex demodulation is the method of choice for calculating time-frequency representations in **BESA** software (Hoechstetter, Bornfleth, Weckesser, Ille, Berg, and Scherg 2004). Complex demodulation is defined by

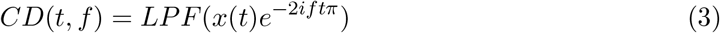

Where *LP F* represents a low-pass filtering operation. The multiplication by *e*^*−*2*iftπ*^ demodulates the signal to baseband, and the low-pass filter removes high-frequency components.

The primary benefit of complex demodulation is its high efficiency. The notable disadvantage of complex demodulation is its scarce representation in the literature, rendering results less comparable to more common TF methods such as STFT or wavelets. This scarcity may also imply a lack of comprehensive understanding of the method’s potential artifacts or biases in the context of MEG/EEG data.

Our implementation of complex demodulation first multiplies the data with a series of sine waves with specified frequencies, followed by applying Butterworth low-pass filters (Butterworth 1930). Users may specify the frequency and order of low-pass filters if desired. Depending on the cut-off frequency and order of the low-pass filter, different temporal and frequency resolution can be achieved. Complex demodulation is calculated using

~~~
>> TF = nf_demodulation(data, Fs, freqs, lowpassF, order, plt);
~~~

Where data is a 1/2/3D tensor of dimensions channels X time X trials, Fs is the sampling rate of the data in Hz, freqs is a vector of center frequencies for the sine waves, lowpassF is the frequency of the low-pass filter in Hz, order is the order of the low-pass filter, and plt is 0 or 1 indicating whether or not the user would like a summary plot to be produced following transformation.

#### Continuous Wavelet Transform

The Continuous Wavelet Transform (CWT) transforms data to a TF representation using wavelets, which are complex sine waves multiplied by a window function that localizes the wavelet energy at the center time (Grossmann and Morlet 1984). The CWT scales and translates a “mother wavelet” by dilating it in time, localizing different frequencies in the process. The transform itself is carried out by convolving the wavelets with the time-domain signal, quantifying the similarity of the time-domain signal to the time-domain wavelet. The CWT is defined by

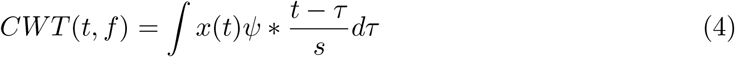

where *ψ* represents the mother wavelet and *s* represents a scale factor equivalent to inverse frequency. The mother wavelet is a complex sine wave multiplied by a Gaussian, defined as

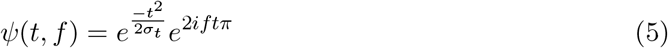

where *σ* is a scaling factor that determines the width of the wavelet.

The CWT has a benefit over the STFT in that it produces a non-uniform tiling of the time-frequency representation. This non-uniform tiling produces higher temporal resolution at higher frequencies and higher frequency resolution at lower frequencies. This feature can be particularly useful in MEG/EEG analysis, as it better accommodates the different timescales over which neural oscillations in different frequency bands tend to occur, correcting for some of the inherent weaknesses of the STFT. The Morlet wavelet, the type of mother wavelet commonly used in MEG/EEG analysis, provides a good balance between time and frequency resolution and has a straightforward interpretation due to its Gaussian shape in both time and frequency domains. Our implementation of the CWT uses the MATLAB `cwt.m` function. CWT is calculated using

>> TF = nf_cwt(data, Fs, plt);

Where data is a 1/2/3D tensor of dimensions channels X time X trials, Fs is the sampling rate of the data in Hz, and plt is 0 or 1 indicating whether or not the user would like a summary plot to be produced following transformation. Note that for Morlet wavelets, the MATLAB function `cwt.m` does not allow modifying the number of cycles.

#### Discretized Continuous Morlet Wavelet Transform

A close relative to the CWT, the discretized continuous wavelet transform (DCWT) adapts the CWT for use with discrete frequency vectors. A key advantage of the DCWT is its computational efficiency, which improves upon that of the CWT. By evaluating the transform at a select vector of frequencies instead of a near-continuous range, the DCWT reduces the computational demands of the transform, which can be a significant advantage when dealing with large MEG/EEG datasets (Cohen 2014). Furthermore, the DCWT allows for greater flexibility by enabling wavelet construction with a user-specified number of cycles at each frequency. This provides the user with control over the time-frequency resolution at each frequency. The discretized continuous wavelet transform is defined by the same equation as the CWT.

However, the benefit of computational efficiency in the DCWT comes with a trade-off in frequency resolution. The frequency resolution is less fine-grained in the DCWT than in the CWT due to the use of a discrete frequency vector. As with the CWT, the Morlet wavelet is the most common wavelet shape used in analyzing M/EEG data with the DCWT.

Our implementation of the DCWT utilizes wavelets that are unit-normalized in the frequency domain, ensuring that the amplitude of the time-frequency representation reflects the actual signal power, regardless of the number of cycles specified (Cohen 2019). This maintains consistency in amplitude measures across different settings of the wavelet width parameter, which is controlled by specifying the number of cycles in the wavelet at each frequency. DCWT is calculated using

~~~
>> TF = nf_wavelet(data, Fs, freqs, cycles, plt);
~~~

Where data is a 1/2/3D tensor of dimensions channels X time X trials, Fs is the sampling rate of the data in Hz, freqs is a vector of center frequencies for the wavelets, cycles is the number of cycles in each wavelet, and plt is 0 or 1 indicating whether or not the user would like a summary plot to be produced following transformation. Cycles can be specified as a single element (e.g. 3) in which case all wavelets will contain 3 cycles, or as a two-element vector (e.g. [3 8]) in which case the number of cycles will being at 3 at the lowest frequency and will linearly increase to 8 cycles at the highest frequency.

#### Stockwell Transform (S-transform)

The Stockwell Transform, or S-transform (Stockwell, Mansinha, and Lowe 1996), is based on the Fourier Transform, specifically starting by taking the Discrete Fourier Transform of the signal. The S-transform then applies a set of frequency-dependent Gaussian window, akin to the scalable windows of wavelets, introducing an adaptive time-frequency resolution. The Gaussian-windowed signals are then brought back to the time domain via the inverse fast Fourier transform. The width of the Gaussian window changes with frequency, which results in higher temporal resolution at higher frequencies and higher frequency resolution at lower frequencies. This is akin to the non-uniform time-frequency resolution provided by wavelet methods, such as the CWT and DCWT. The S-transform is defined by

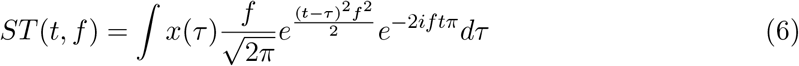

The S-transform combines the advantages of both STFT and wavelet methods, and can potentially provide a clearer, more detailed time-frequency representation of M/EEG data compared to either method used alone. However, the S-transform is more computationally expensive than either STFT or wavelets due to its adaptability in time-frequency resolution and the inverse Fourier Transform step.

The S-transform is calculated using

~~~
>> TF = nf_stransform(data, Fs, factor, plt);
~~~

Where data is a 1/2/3D tensor of dimensions channels X time X trials, Fs is the sampling rate of the data in Hz, factor controls the resolution of the transform (generally 1 but can be 3 for higher resolution), and plt is 0 or 1 indicating whether or not the user would like a summary plot to be produced following transformation.

### 3.3. Quadratic Time-Frequency Decomposition Methods

Quadratic time-frequency methods can provide high-resolution information in both time and frequency simultaneously, making them particularly suited for complex signals that exhibit rapid and non-stationary changes in frequency content. However, this comes at the cost of increased computational complexity and cross-term interference, which can result in time-frequency representations that are more challenging to interpret.

#### Wigner-Ville Distribution

The Wigner-Ville distribution (WVD) is a quadratic time-frequency representation that was originally developed in quantum mechanics (Wigner 1932), but has found application in many areas of signal processing. It is infrequently used in M/EEG analysis primarily due to the presence of “cross-terms” that complicate interpretation of the resulting time-frequency representation. To introduce notation for the WVD first define the local (instantaneous) autocorrelation function, which reflects a signal back on itself to assess self-similarity at different time lag values, as:

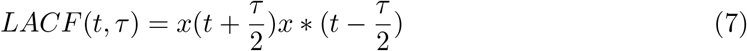

The WVD then is computed as the Fourier transform of *LACF* (*t, τ*) with respect to *τ* :

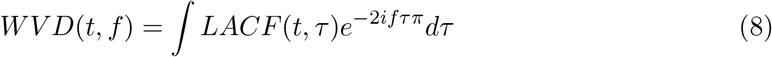

This produces a time-frequency representation that has the highest mathematically possible resolution, as the resolution is not constrained by a fixed window length or shape as in the case of STFT, nor by the number of wavelet cycles as in the case of wavelet-based methods. This can, in theory, offer very detailed insight into the time-frequency structure of a signal. However, this advantage is tempered by a significant disadvantage: the aforementioned cross-terms. Cross-terms are interference patterns that occur between different components of a multi-component signal. These cross-terms can produce spurious peaks in the time-frequency representation that do not correspond to any real feature of the signal, but are instead artifacts of the computation process. This can render the WVD very difficult to interpret, especially for complex signals such as M/EEG data, which consist of multiple overlapping oscillatory components. **NeuroFreq** does not implement the WVD due to its inherent issues with cross-terms. However, the concepts underlying the WVD inform the development and application of other quadratic methods that are included in the toolbox. These methods modify the WVD to suppress the cross-terms and provide more interpretable time-frequency maps.

#### Binomial Reduced Interference Distribution

Cohen’s class of time-frequency transforms (Cohen 1995) encompasses a broad range of methods that apply a kernel to modify the WVD. These kernels are best understood in terms of the ambiguity function *AF* (*τ, σ*), obtained by calculating *LACF* (*t, τ*) and then taking the Fourier transform with respect to *t*, instead of *τ* as used in the WVD:

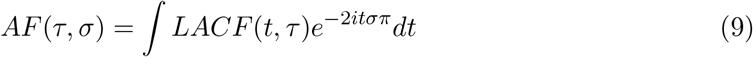

The ambiguity function shifts the signal auto-terms to the origin while concentrating the cross-terms away from the origin. Thus, application of a kernel surrounding the origin in the ambiguity domain serves to suppress the cross-terms inherent in the WVD while retaining the auto-terms. After applying the kernel, the masked ambiguity function can be transformed back to the time-frequency domain by applying two Fourier transforms.

The Binomial Reduced Interference Distribution (BRID) (Jeong and Williams 1992) adopts a cross-shaped kernel in the ambiguity domain. The binomial kernel downweights the cross-terms that typically occur away from the origin. The binomial kernel is defined as

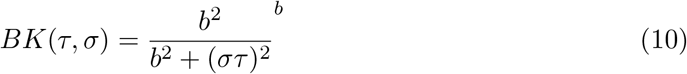

where *b* is a parameter controlling the order of the binomial kernel, usually set to 2. While the BRID is less computationally efficient than linear TF methods, it can produce TF transforms with higher clarity in some cases. The BRID attenuates cross-terms but does not eliminate them, and has lower resolution than the WVD due to the smoothing kernel. The toolbox uses inline code released under a GNU Public License to compute the binomial kernel RID. Since the binomial RID returns a real-valued output, the result has a ‘power ‘ field but no ‘phase ‘ field. The binomial RID is computed using

~~~
>> TF = nf_ridbinomial(data, Fs, fRes, maxLags, makePos, plt);
~~~

where data is a 1/2/3D tensor of dimensions channels X time X trials, Fs is the sampling rate of the data in Hz, fRes is the desired frequency resolution of the output in Hz (e.g. 0.5 produces a TF representation with frequency steps equal to 0.5 Hz), maxLags is the length of the autocorrelation window in seconds, makePos is 0 or 1 indicating whether the distribution should return only positive energy, and plt is 0 or 1 indicating whether or not the user would like a summary plot to be produced following transformation. Shorter autocorrelation windows more effectively mask cross-terms, but also reduce the TF resolution. Note that only interference terms can take on negative values, so makePos can potentially return a clearer TF representation.

#### Born-Jordan Distribution

The Born-Jordan Distribution (BJD) is a quadratic time-frequency distribution in Cohen’s class. The BJD was originally derived in (Cohen 1966), and its general properties were studied by (Jeong and Williams 1992). The kernel of the BJD is a rectangular function in the time-lag domain, which completely removes the cross-terms in the absence of noise. This makes the BJD particularly well suited to the analysis of multi-component signals, where cross-terms can otherwise introduce substantial interference. In practice, the cross-term-free property is slightly compromised due to the presence of noise and other non-idealities in real-world data. However, even in such cases, the BJD often outperforms other members of Cohen’s class in terms of cross-term suppression. The kernel of the BJD is defined as:

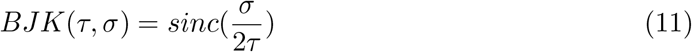

Much like the binomial RID, the BJD is less computationally efficient than linear TF methods, but can produce TF transforms with much higher clarity in some cases. In the presence of noise the BJD attenuates cross-terms but does not eliminate them, which is critical to keep in mind since M/EEG data always contain some level of noise. Our toolbox uses inline code released under a GNU Public License to compute the Born-Jordan distribution. Since the Born-Jordan distribution returns a real-valued output, the result has a ‘power ‘ field but no ‘phase ‘ field. The BJD is computed using

~~~
>> TF = nf_ridbornjordan(data, Fs, fRes, maxLags, makePos, plt);
~~~

Where data is a 1/2/3D tensor of dimensions channels X time X trials, Fs is the sampling rate of the data in Hz, fRes is the desired frequency resolution of the output in Hz (e.g. 0.5 produces a TF representation with frequency steps equal to 0.5 Hz), maxLags is the length of the autocorrelation window in seconds, makePos is 0 or 1 indicating whether the distribution should return only positive energy, and plt is 0 or 1 indicating whether or not the user would like a summary plot to be produced following transformation. Shorter autocorrelation windows more effectively mask cross-terms, but also smear the TF representation. Note that only interference terms can take on negative values, so makePos can potentially return a clearer TF representation.

#### Reduced Interference Distribution Rihaczek

The Rihaczek Complex Energy Spectrum (RCES) (Rihaczek 1968) incorporates both the time shifts *τ* and frequency shifts *σ* in its phase function, unlike other quadratic time-frequency distributions which only consider frequency shifts. This results in the RCES uniquely providing simultaneous energy and phase distribution information. The Rihaczek distribution can be defined as a kernel applied to the WVD in the ambiguity domain, where the Rihaczek kernel is defined as

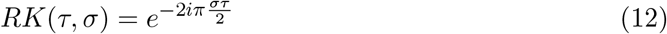

However, like the WVD, the RCES is susceptible to cross-term interference which complicates interpretation of its time-frequency representation. To address this, the Reduced Interference Distribution Rihaczek (RID-Rihaczek) was proposed (Aviyente, Bernat, Evans, and Sponheim 2011), combining the power of the RCES with the reduced interference kernel approach of Cohen’s class. The RID-Rihaczek modifies the Rihaczek CES by introducing a Choi-Williams kernel (Choi and Williams 1989), an exponential kernel that attenuates the contributions of cross-terms. The Choi-Williams kernel is defined as:

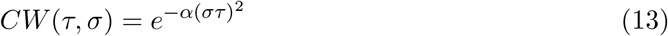

where *α* is a parameter that controls the suppression of cross-terms. This kernel is designed to mitigate cross-term interference while preserving the distinctive phase information that the RCES offers. In essence, the RID-Rihaczek represents a balanced trade-off between the preservation of energy-phase information and the minimization of cross-term interference.

The toolbox utilizes inline code released under a GNU Public Use License to compute the RID-Rihaczek. The ‘power ‘ field returns the absolute value of the real part of the Rihaczek spectrum, since only the real part contributes to the marginals. The ‘phase ‘ field returns the phase angle of the complex result. The RID-Rihaczek is computed using

~~~
>> TF = nf_ridrihaczek(data, Fs, fRes, cwkernel, makePos, plt);
~~~

Where data is a 1/2/3D tensor of dimensions channels X time X trials, Fs is the sampling rate of the data in Hz, fRes is the desired frequency resolution of the output in Hz (e.g. 0.5 produces a TF representation with frequency steps equal to 0.5 Hz), cwkernel controls the size of the Choi-Williams kernel, makePos is 0 or 1 indicating whether the distribution should return only positive energy, and plt is 0 or 1 indicating whether or not the user would like a summary plot to be produced following transformation. Larger kernels more effectively mask cross-terms, but also smear the TF representation.

## 4. Demonstrations

### 4.1. Synthetic Data

Here we demonstrate the TF algorithms included in **NeuroFreq** using synthetic data generated by (Arts and van den Broek 2022) for benchmarking time-frequency algorithms. The two synthetic datasets are composed of multiple time-varying signals sampled at 500 Hz. These synthetic signals are described by 1) three 5-s sine waves with frequencies that gradually oscillate between 5-6 Hz, 20-22 Hz, and 100-110 Hz, fluctuating with a periodicity of 1 Hz;2)two 5-s sine waves with frequencies that change linearly from 5-50 and 100-50 Hz; and 3) three 10-s sine waves with frequencies of 0.5, 1, and 2 Hz. The three sets of signals are separated by 0.5 s and are multiplied by a Gaussian window function to mitigate discontinuities at the boundaries, as described in (Arts and van den Broek 2022). One of the synthetic datasets contained only the aforementioned sine waves (that is, clean data), and the other was combined with Gaussian noise that produced a signal-to-noise ratio of 1. Both datasets have a total duration of 21.0 s.

The algorithms were run with the following parameters: STFT (brief window): window = 0.5 s, overlap = 80%, frequency resolution = 0.1 Hz; STFT (long window): window = 5 s, overlap = 95%, frequency resolution = 0.1 Hz; filter-Hilbert: filter bandwidth = 0.25 Hz, filter order = 3; Complex demodulation: low-pass filter frequency = 2 Hz, filter order = 3; discretized continuous wavelet transform: cycles = [3 8]; S-transform: factor = 1. CWT does not have user-selectable parameters. The quadratic algorithms were run with default parameters; specifically, all algorithms returned n(times) frequency points, and the binomial RID and Born-Jordan distribution used autocorrelation windows with length 2*n(times) while the RID-Rihaczek used a kernel parameter of 0.001.

Figure 2 depicts the output of each of the linear TF decompositions on synthetic data. Outputs for the clean (uncorrupted) data are shown in the left column, and outputs for noisy (corrupted) data are shown in the right column. Callouts zoom into the 0-3 Hz and 15-20 s segment to show resolution for low-frequency sine waves. Several salient points were noted. First, the STFT has extremely high noise-canceling properties which comes at the expense of fixed TF window resolution. The short window does not allow separation of very low frequency waves, but the longer window obscures high-frequency dynamics. Demodulation also has very high clarity for the primary signal components but shows large edge effects which likely arise from the multiplication of untapered sine waves. Demodulation also has poor resolution for the low-frequency rhythms. Filter-Hilbert and DCWT showed very similar output for both clean and noisy data. CWT and S-transform were the only transforms with sufficient resolution at low frequencies to observe all three separate low-frequency oscillations.

**Figure 2.**
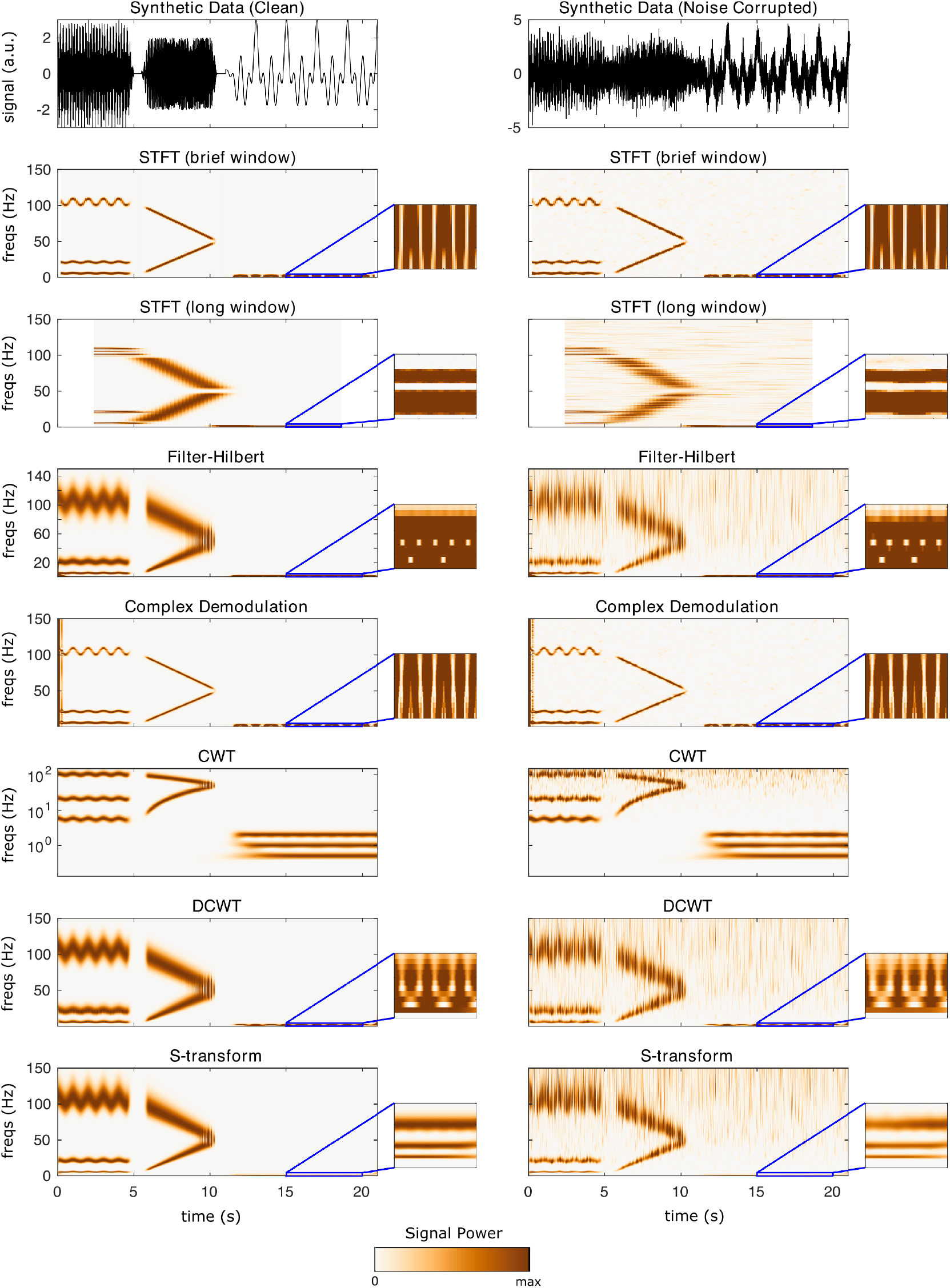
Results of linear TF transformations on synthetic data. Left column is for an uncorrupted time-domain signal, right column is the output for the same signal corrupted with noise. Callouts show the result zoomed into 0-3 Hz and 15-20 seconds, to visualize resolution for the 0.5, 1, and 2-Hz waves. Note that CWT is shown on a logarithmic scale; this logarithmic frequency spacing is part of the CWT’s definition.

Figure 3 depicts the output the quadratic TF distributions on synthetic data. The Wigner-Ville distribution famously suffers from cross-term contamination in the presence of multi-component signals. These cross-terms are evident as oscillating spectral content in between signal auto-terms. The binomial RID and the Born-Jordan distribution improve cross-term contamination, but do not eliminate it. In the case of the binomial RID, oscillating spectral content is present between 20-100 Hz from 0-5 s as a cross-term originating from the 20 & 100 Hz auto-terms, and oscillating cross-terms are also visible around 50 Hz from 6-10 s originating from the two chirp auto-terms. Cross-term interferences are also visible in high frequencies from 12-21 s, originating from interferences between the very low-frequency auto-terms. These interferences are still present, although reduced, in the Born-Jordan distribution. The RID-Rihaczek shows lower clarity for signal auto-terms than the binomial RID or Born-Jordan distribution, and is degraded significantly in the presence of noise. Each of the quadratic distributions was able to resolve the three separate low-frequency waves, but with substantial interferences. The primary conclusion is that the quadratic TF distributions have high uniform resolution and show high clarity, but are susceptible to noise, as the clarity of the outputs degrades significantly for contaminated signals.

**Figure 3.**
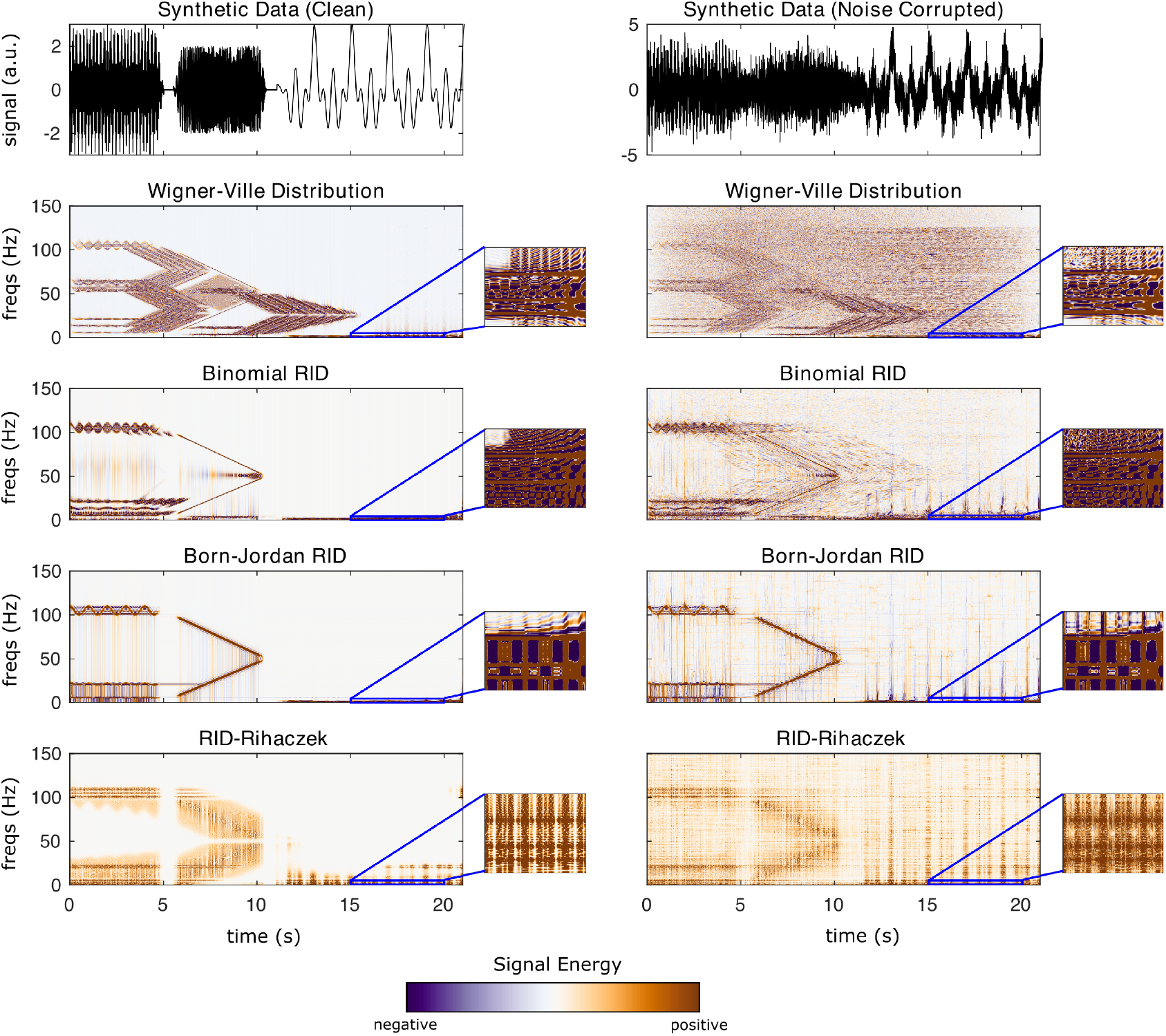
Results of quadratic TF transformations on synthetic data. Left column is for an uncorrupted time-domain signal, right column is the output for the same signal corrupted with noise. Callouts show the result zoomed into 0-3 Hz and 15-20 seconds, to visualize resolution for the 0.5, 1, and 2-Hz waves.

### 4.2. Experimental EEG Data

This section demonstrates the different TF algorithms included in **NeuroFreq** using experimental EEG data. These EEG data are publicly released as tutorial data with **EEGLAB**, and consist of 32 channels of EEG sampled at 128 Hz while a subject performed 80 trials of a visual cognitive task. Following TF computation, data were averaged over trials using `nf_avebase.m`, which averaged but did not baseline correct TF power, and which averaged phase over trials to get ITPC estimates. These data show several common spectral phenomena observed in experimental EEG data, including strong alpha oscillations over parietal sensors and post-stimulus phase locking of low frequencies, making these data suitable for demonstration of the TF decomposition algorithms included in **NeuroFreq**.

The algorithms were run with the following parameters: STFT: window = 0.5 s, overlap = 80%; filter-Hilbert: filter bandwidth = 1 Hz, filter order = 3; Complex demodulation: low-pass filter frequency = 2 Hz, filter order = 3; discretized continuous wavelet transform: cycles = [3 8]. CWT and S-transform do not have user-selectable parameters. The quadratic algorithms were run with the following parameters: binomial RID & Born-Jordan: resolution = 0.5 Hz, kernel = 2*n(times), makePos = 0; RID-Rihaczek: resolution = 0.5 Hz, kernel = 0.001.

Figure 4 displays TF surfaces at sensor Pz, and scalp topographies averaged over all time points and frequencies from 9-14 Hz. All TF decomposition methods reveal strong alpha power centered at 10 Hz, with a scalp focus at parietal/occipital sites. Some TF methods also indicate an increase in low-frequency power from 200-500 ms, most notably STFT, filter-Hilbert, demodulation, binomial RID, and Born-Jordan. This dynamic increase in low-frequency power is not observed as strongly for the CWT and S-transform. The quadratic TF distributions show a more narrow alpha-band synchronization than the linear methods, which is expected given the high resolution of these methods.

**Figure 4.**
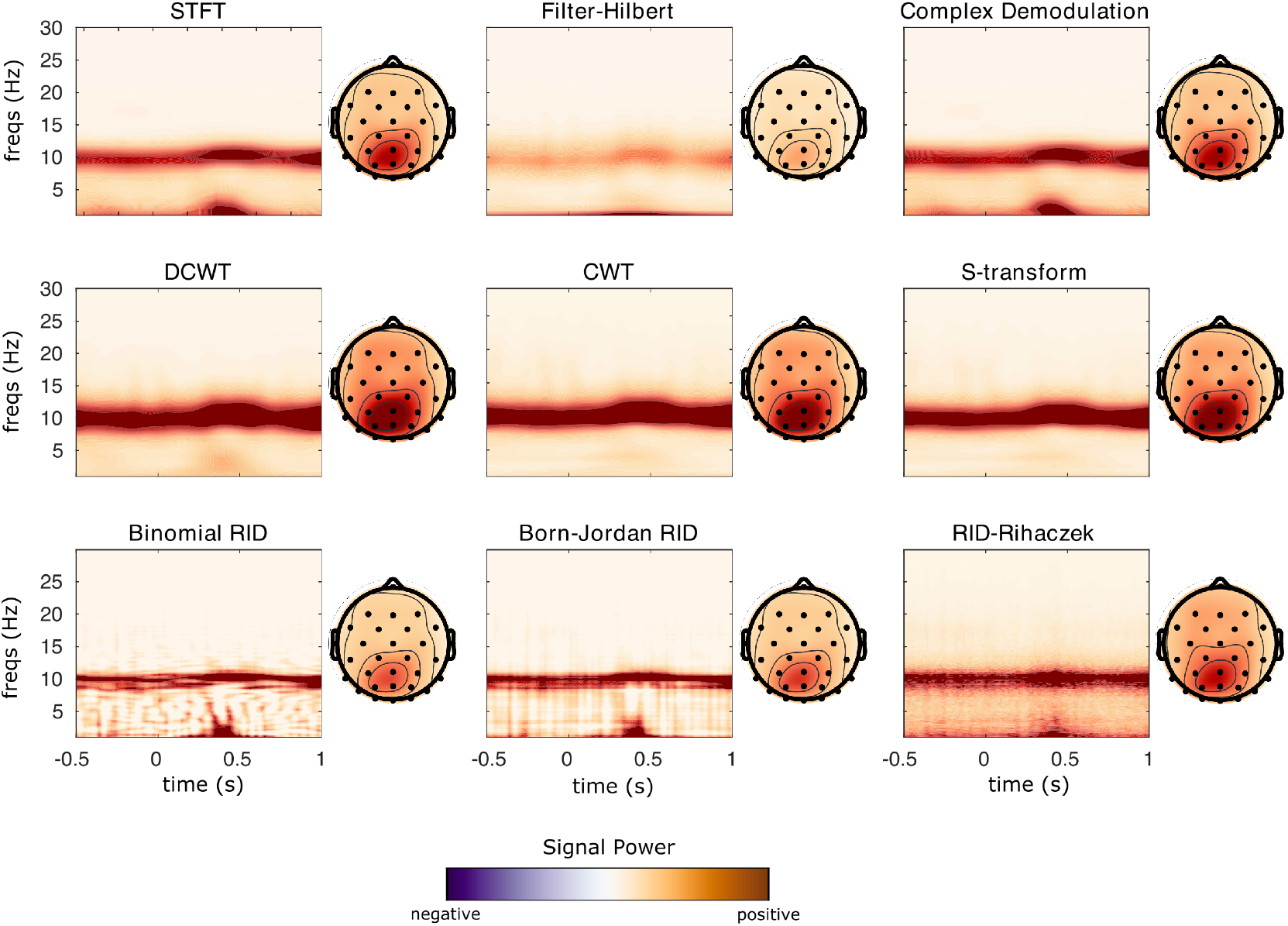
Power results from each TF transformation applied to sample EEG data from a visual experiment. TF transforms were calculated in single trials and results were averaged over 80 trials for plotting. TF surfaces are displayed at Channel Pz, where the participant showed strong alpha rhythm synchronization.

Figure 5 displays ITPC surfaces at sensor Fz, and scalp topographies averaged over all time points and frequencies below 5 Hz. Again, all TF decompositions show similar post-stimulus low-frequency phase locking, with a scalp focus at anterior right-central sensors. The localization of this phase-locking in time differs markedly between the different TF methods, with the tightest temporal localization observed for STFT, demodulation, and RID-Rihaczek methods and comparatively more smeared time localizations for CWT, S-transform, DCWT, and filter-Hilbert methods.

**Figure 5.**
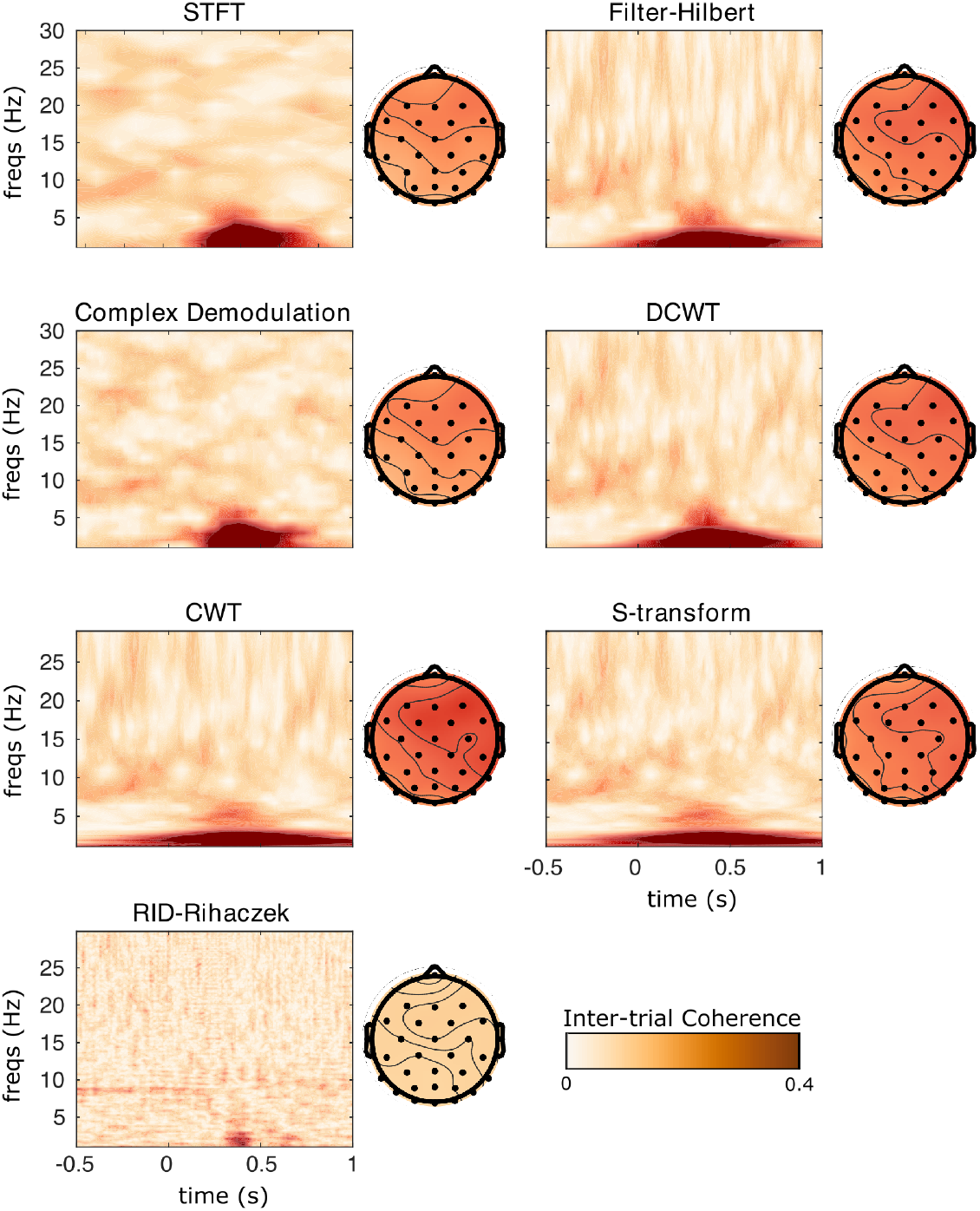
Inter-trial phase coherence results for each TF transformation applied to sample EEG data from a visual experiment. TF transforms were calculated in single trials and results were averaged over 80 trials to calculate ITPC. Surfaces are displayed at Channel Fz, where the subject showed strong delta phase synchronization.

## 5. Summary and discussion

This paper describes the NeuroFreq toolbox for calculating time-frequency decompositions using MATLAB. The toolbox includes a wider variety of linear and quadratic TF decompositions than current packages, and is optimized for M/EEG data structures with support for data structures potentially including data from multiple channels and trials. While the consistent TF transforms themselves are the primary contribution of the toolbox, the toolbox also includes utilities for averaging, baseline correction, and viewing the calculated TF structures.

The toolbox uses a consistent structure for outputting the results of TF transformation, and can accept as input either data matrices or **EEGLAB** .set structures. The **NeuroFreq** toolbox makes flexible and consistent TF analysis available to a wider range of researchers.

## Acknowledgements

I am especially thankful to Dr. Olivia Calvin and Dr. Erich Kummerfeld for helpful comments on this manuscript. This toolbox was developed and tested over an extended period of time. Over this time I was supported by the National Institutes of Health’s National Center for Advancing Translational Sciences, grants TL1TR002493 and UL1TR002494, and the National Institute of Mental Health, grant T32-MH115866. The content is solely my responsibility and does not necessarily represent the official views of the National Center for Advancing Translational Sciences or the National Institute of Mental Health. I used many excellent public code resources in developing the NeuroFreq toolbox. These include: 1) utilities by M.X. Cohen for wavelet convolution and TF visualization, publically available from https://github.com/mikexcohen, 2) utilities by J. O ‘Neill for discrete time-frequency distributions, publicly available from https://tfd.sourceforge.net/, 3) utilities by S. Aviyente for RID-Rihaczek computation, available at https://github.com/SPLab-aviyente, 4) an original implementation of the S-transform by R. Stockwell, available at https://www.mathworks.com/matlabcentral/fileexchange/51808-time-frequency-distribution-of-a-signal-using-s-transform-stockwell-transform.

## Code and Documentation

**NeuroFreq** is hosted on Github at https://github.com/erawls-neuro/NeuroFreq_public.

**NeuroFreq**’s documentation can be found at https://neurofreq-public.readthedocs.io/en/latest/index.html.

## Preprint Disclaimer

This is an early version of the paper describing the **NeuroFreq** Toolbox. This document, and the utilities described here, are likely to change substantially before peer-reviewed publication. Please with any suggestions or for any additional information or questions.

